# Late-life fitness gains explain the absence of a selection shadow in ants

**DOI:** 10.1101/2021.02.24.432699

**Authors:** Luisa M. Jaimes Nino, Jürgen Heinze, Jan Oettler

## Abstract

A key hypothesis for the occurrence of senescence is a decrease in the selection strength because of low late-life fitness – the so-called selection shadow. However, in social insects, aging is considered a plastic trait and senescence seems to be absent. By life-long tracking of 102 ant colonies, we find that queens increase the production of sexuals in late life regardless of their absolute lifespan or worker investment. This indicates a genetically accommodated adaptive shift towards increasingly queen-biased caste ratios over the course of a queens’ life. Furthermore, mortality decreased with age, supporting the hypothesis that aging is adaptive. We argue that selection for late life reproduction diminishes the selection shadow of old age and leads to the apparent absence of senescence in ants, in contrast to most iteroparous species.

**Significance Statement:** Social insects are extraordinary with regard to the absolute age of queens of some species as well as the age differences between queens and short-lived workers. Yet, ultimate causes explaining these aging patterns remain poorly understood. By manipulating the investment ratio into queens and workers we studied the effect on lifespan and temporal reproductive investment of queens. We show that queens shift to the production of sexuals late in life, independent of social context (colony size) or individual quality (reproductive output and lifespan). Such late fitness gains result in maintenance in the selection strength with age, and thus explains the absence of senescence in queens.

## Introduction

The near-ubiquitous occurrence of senescence, a trait without any apparent fitness benefit, has been explained by two classic prevailing theories, mutation accumulation (MA) and antagonistic pleiotropy (AP) (Haldane 1941; Kirkwood 1977; Hughes and Reynolds 2005; Maklakov et al. 2015; Flatt and Partridge 2018). These theories rely on the basic assumption of the existence of a “selection shadow”. Selection against age-specific mortality decreases with age, and is proportional to the number of offspring that come from parents that survived to that age (Hamilton 1966; Moorad et al. 2020). This in turn can lead to loss-of-function and senescence. Therefore, mutations with an effect late in life could accumulate (MA) and/or genes with beneficial effect early in life can be selected for even if they are detrimental to the organism later in life (AP) (Maklakov and Chapman 2019). Such antagonistic pleiotropy is visible in the conflict over resource allocation to either reproduction or somatic maintenance (i.e. disposable soma theory, or energy trade-offs) (Kirkwood 1977; Kirkwood and Austad 2000) and/or in suboptimal gene regulation after maturation (functional trade-offs) (López-Otín et al. 2013; Maklakov and Chapman 2019). However, the huge diversity of aging patterns across metazoan species (Jones et al. 2014; Cohen 2018) calls for a critical evaluation of the general validity of these theories.

When considering social insect reproductives (e.g. ant queens), the basic assumptions underlying MA and AP may not be met (Monroy Kuhn and Korb 2016; Toth et al. 2016; Lucas and Keller 2017). First, the costs of reproduction (energy intake, brood care) are outsourced to the workers. Second, it is posited that selection at high ages is reinforced if there is no marked decrease in fecundity after maturity (Keller and Genoud 1997), predicting that classic energy trade-offs may be shaped differently or may not even exist. This has found support in studies of the model ant *Cardiocondyla obscurior* (Schrempf et al. 2005; Schrempf et al. 2011; Oettler and Schrempf 2016) where queens exhibit a positive correlation between lifespan and reproduction (Kramer et al. 2015). Furthermore, experimental manipulation of the fecundity of *C. obscurior* queens does not affect lifespan (Schrempf et al. 2017) and queens appear to have only a very brief period of senescence (Heinze and Schrempf 2012). Additional evidence for the lack of senescence comes from a gene expression study, which showed a lack of transcriptional senescence in old *C. obscurior* queens, possibly facilitated by strong selection at old age and by well-regulated anti-senescence mechanisms (Harrison et al. 2021). So how did this exceptional pattern of positive correlation between fecundity and lifespan, and the absence of senescence, evolve, and are there trade-offs at the colony level that shape queen lifespan instead?

In order to study aging of social insect queens it is necessary to consider the colony as a superorganism (Boomsma and Gawne 2018), comprising a soma- (i.e. workers) and a germline (i.e. queens and males), where the investment into both castes is related and affects overall fitness (Bourke 2007; Kramer and Schaible 2013). Therefore, by manipulating the investment into caste ratio, we expected to find trade-offs at the level of the queen (i.e. lifespan and productivity of queens) and/or at the level of the colony (investment in queen/worker offspring), if they exist. To this end, we monitored the lifetime production of 102 individual queens in colonies, which were standardized weekly to 10, 20 or 30 workers (Fig. S1A and Fig. S1B), corresponding to the natural colony size variation (Schrader et al. 2014, Fig. S2). In order to compare queen mortality with worker aging patterns, we tracked the survival of 40 workers kept in colonies with 10 or 20 marked nestmate workers.

## Results and Discussion

The treatment did not affect total egg production (Fig. 1A, 10 vs. 20 workers: z-value = −0.38, p = 0.70 and 10 vs. 30: z-value = −0.96, p = 0.34) or worker pupae production (Fig 1B, 10 vs. 20 workers: z-value = 0.09, p = 0.93 and 10 vs. 30: z-value = −0.39, p = 0.70).The treatment did not affect lifespan (Fig. S3A, Cox proportional hazard regression model, Likelihood ratio test, X^2^=1.57, p = 0.46), which was highly variable across treatments (Variation coefficient: 32.2%, Fig. S3B). This confirms a previous study that did not find a causal correlation between lifespan and fecundity (Schrempf et al. 2017).

**Fig. 1.**
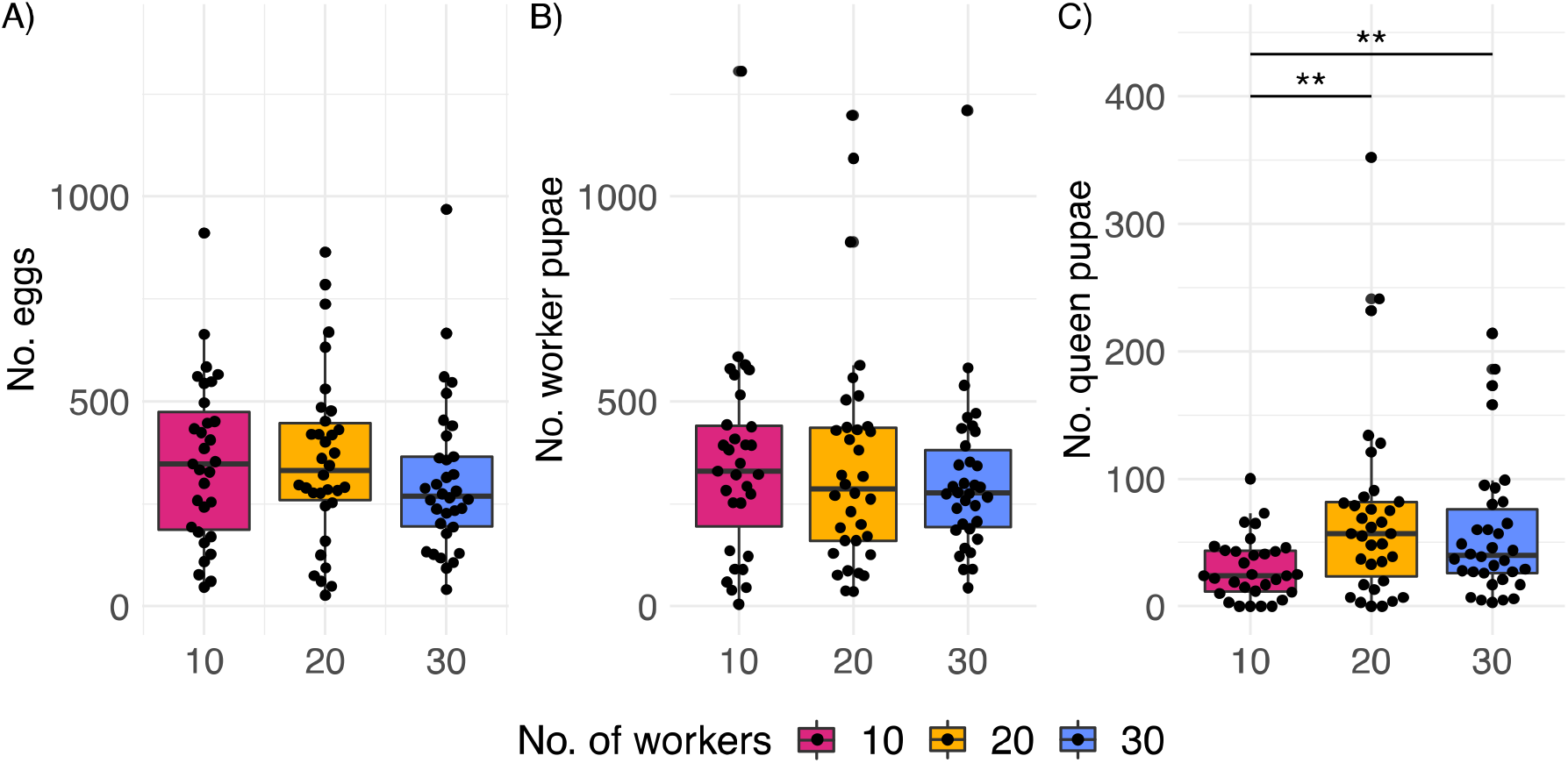
Productivity of *C. obscurior* colonies across treatments, A) Total number eggs, B) 157 worker pupae, and C) of queen pupae. Significant differences are given with ** for p<0.01 158 and *** for p<0.001.

We hypothesized that queens which experienced a worker shortage would compensate by investing less into queen production. This assumes queen control over caste fate in *C. obscurior*, in contrast to other species where caste is environmentally or genetically determined (Corona et al. 2016). Indeed, queens with 10 workers (n=31) produced significantly fewer queen pupae than queens with 20 (n=34) (glmmTMB z-value = 2.81, IRR = 1.97, p = 0.005) and 30 workers (n=34) (z-value = 2.58, IRR = 1.78, p = 0.009, Fig. 1C) with no significant differences between 20 and 30 workers (z-value = −0.49, p = 0.877).

A first peak in queen-biased investment occurs at an earlier age (Fig 2A), followed by an increasing queen bias with increasing age (Fig 2B). Importantly, this caste ratio shift is independent of queen lifespan (queens with a lifespan below and above the mean lifespan of 25 weeks, Fig. 2C and D respectively), implying that queens are selected for late life investment into queens. Founder queens invest first in growing numbers of workers (ergonomic phase) and subsequently in the production of new sexuals, when the colony has reached the threshold required to enter the reproductive phase (Macevicz and Oster 1976; Oster and Wilson 1978; Beekman et al. 1998). Our data suggest that this switch in caste allocation is a fixed trait, independent of colony size and absolute lifespan of queens.

**Fig. 2.**
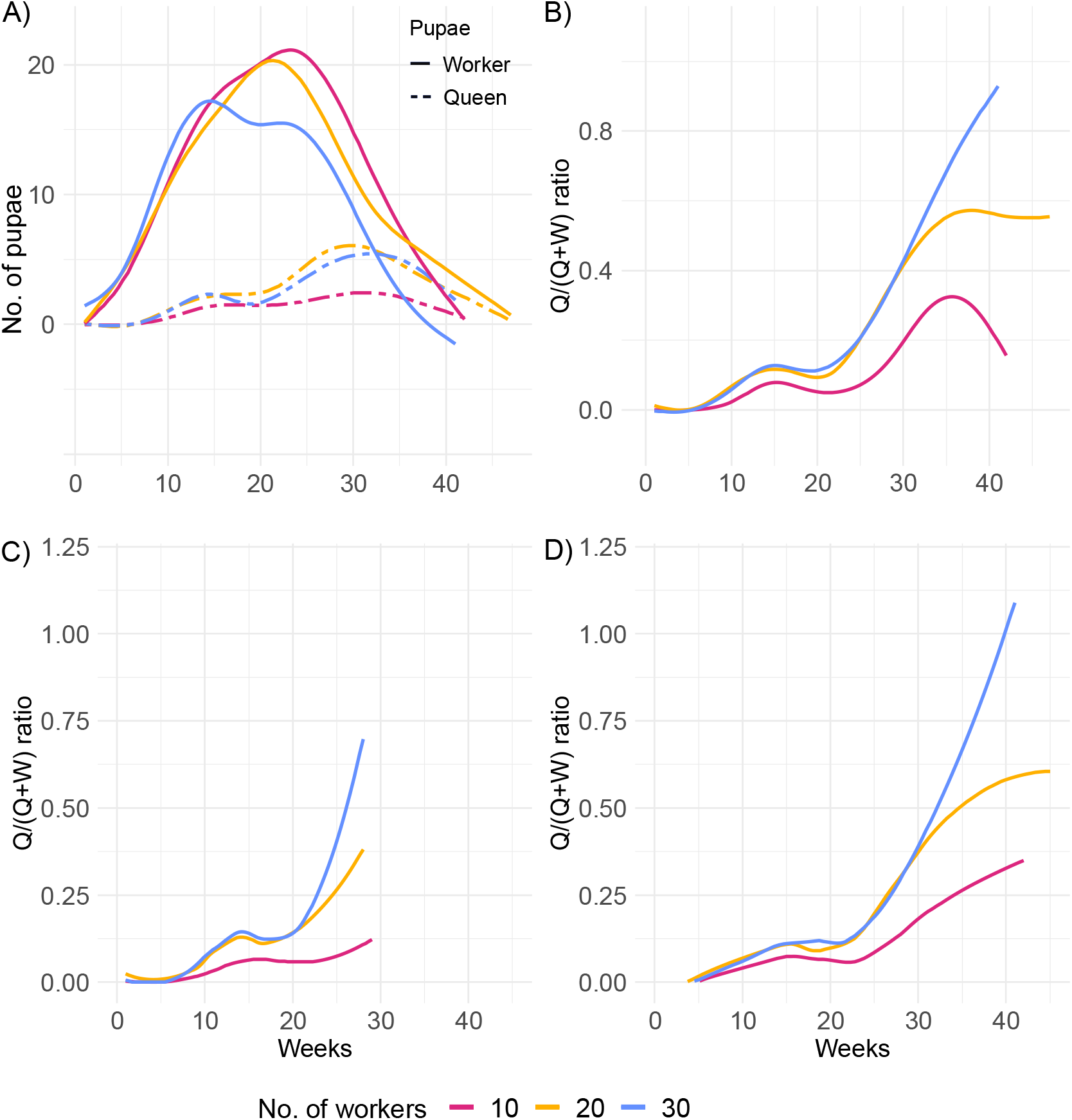
Life-time caste allocation. A) Numbers of worker and queen pupae produced over time, B) Queen/(Queen + Worker) pupae ratio produced by queens, C) Caste ratio for queens with lifespan below (n = 44), and D) above the mean of 25 weeks (n = 55). After queen death, eggs and larvae were allowed to develop into pupae for a final count. Therefore, smooth splines extend ca. 4 weeks after queen death.

In addition to the effect on the caste ratio, the treatment had an effect at the colony level. We explored whether the quality of workers was affected by measuring the head width of workers produced over months 3 to 6 of the queen’s lifetime (~5 workers per month). Head width of workers was 2% and 3% significantly smaller in small colonies than in colonies with 20 (lme F_6,76_ = 2.36, p = 0.026) and 30 workers, respectively (lme F_6,76_ = 3.52, p < 0.002, Fig. 3). This suggests that small colonies are limited, either because the work force is not sufficient to forage or to nurse the brood.

**Fig. 3.**
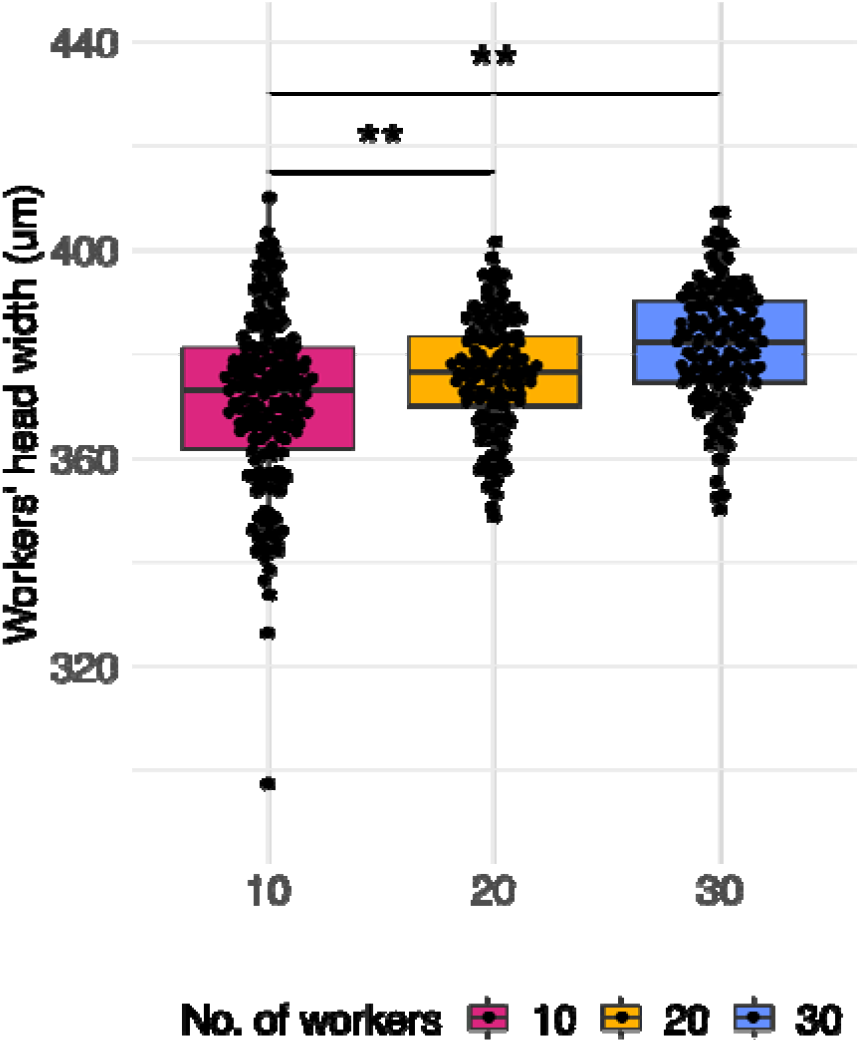
Worker quality per treatment. Head width measurements of workers produced in queens of colonies with 10, 20 and 30 workers. Significant differences are given with * for p < 0.05 and ** for p < 0.01.

To investigate mortality and fecundity patterns we mean-standardized queen age-specific mortality and fecundity (Jones et al. 2014). After an earlier increase in relative mortality, *C. obscurior* ant queens exhibit a decrease below the average level of adult mortality late in life (Fig. 4A), indicating maintenance of selection. Worker production tightly matched the curve of egg production (Fig 4A). Furthermore, in contrast to the prediction that there is no marked decrease in queen fecundity after maturity (Keller and Genoud 1997), relative fecundity reaches a maximum (~16 weeks) before the median lifespan (~26 weeks). However, the relative investment in queen pupae reaches a maximum late in life (~28 weeks). This pattern is not due to the delay in development from egg to pupa, because queen development only lasts ~5 weeks (Schrempf and Heinze 2006). Note that regardless of the differences in time scale, the mean-standardized mortality of workers is very similar to the queen’s mortality, where the relative mortality first increases then decreases (Fig 4B). This suggests that aging is an is a genetically fixed trait expressed by queens and workers alike.

**Fig. 4.**
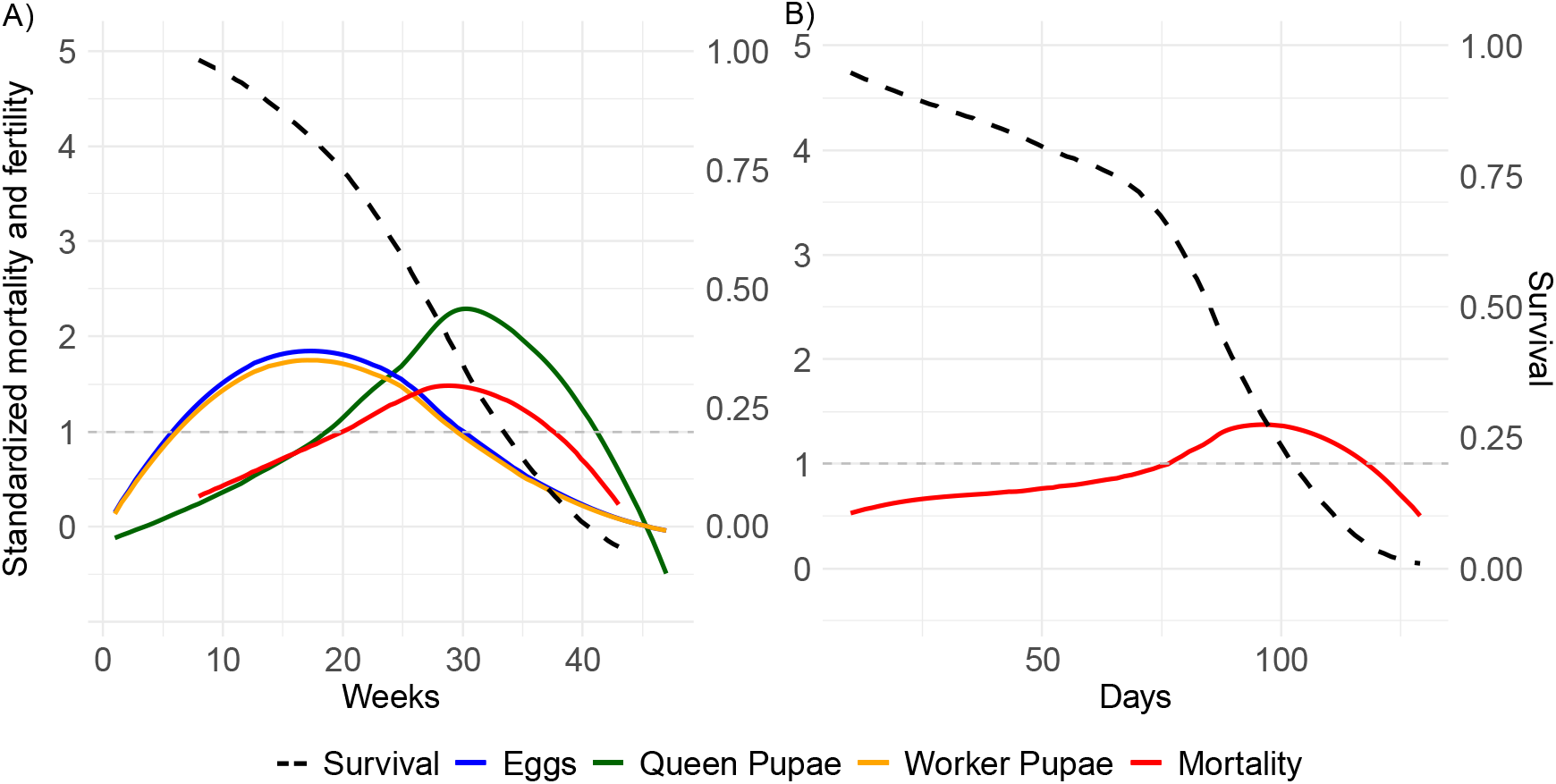
Relative mortality and fecundity as a function of age. Mean-standardization of age by dividing age-specific mortality and fecundity by their mean after maturation (Jones et al. 2014) in A) the queen, and B) worker. The graph uses a Loess smoothing method (span = 0.75) and a confidence interval of 95%. The dashed line at y = 1, indicates when relative mortality and fertility are equivalent to mean mortality and fertility.

## Conclusion

We define senescence as an increase in relative mortality and a decrease in relative fecundity with age, due to a decrease in the strength of selection. Strikingly, *C. obscurior* queens exhibit a decrease in relative mortality with age. Further, while fecundity decreases, investment into queen pupae reaches a maximum late in life, regardless of individual fitness. This indicates that *C. obscurior* queens continue to experience strong selection at high ages. In support of our findings, a study of genes highly expressed in old compared to young *C. obscurior* queens revealed strong purifying selection at a late age (Harrison et al, 2021). In turn, the short selection shadow might have led to the absence of a distinguishable senescence phase. Whether this holds true for ants in general, and whether this drives the evolution of the extraordinary lifespans found in social insects, remains to be studied.

## Acknowledgements

This study was funded by the Deutsche Forschungsgemeinschaft (OE549/2-2). We thank Vera Ermer, Benjamin Dofka, Julia Haschlar, Judith Weber and Lena-Marie Süß for help with the experiment, and Eva Schultner, Tomer Czaczkes and Boris Kramer for comments.

## Author contributions

JH and JO designed the study, LJ and JO produced the data, LJ analyzed the data, LJ and JO wrote the original draft, all authors reviewed and edited the final version of the draft.

## Data and materials availability

All data is available in the main text or the supplementary materials.

## Materials and Methods

We set up 102 freshly eclosed queens from stock colonies of a Japanese population (OypB, from the Oonoyama Park in Naha, Okinawa) established in the laboratory since 2011. The experiment took place between January 2019 and January 2020. Queens were allowed to mate with a single wingless male and were placed in nest boxes with either 10, 20 or 30 workers from the maternal colony (N = 34). This number of workers represents the naturally occurring number in the field and corresponds closely to the first, median and third quantile of number of workers of this population (N = 62, median = 28.5, Fig. S2). The colony was set up with half of the workers selected from inside of the nest near the brood (younger nurses) of the stock colonies, and the other half from outside the nest (older foragers) in order to minimize a putative effect of worker age on the queen (Giehr et al. 2017). Colonies were kept under a 12 h of dark 22°C/12 h light 26°C cycle and fed *ad libitum* three times per week with diluted honey (0.6:1 honey: distillated water), cockroaches and flies. Once per week workers, eggs, and all pupae (worker, queen, winged and wingless male) were counted. Additionally, the number of workers was standardized to the assigned treatment, and newly produced reproductive pupae produced were removed. *C. obscurior* workers are sterile, and all produced offspring originated from the focal queen. Queen control over caste fate was assumed, as caste fate can be determined as early as the last embryonic stage. The number of counted eggs correlates with the production of workers, queens, and the workers and queens together (Fig. S4.A-C, Kendall’s Tau correlation test, p < 0.001: eggs-worker pupae= 0.59, eggs-queen pupae= 0.70, eggs-worker and queen pupae=0.73). Pupae might have been counted more precisely than eggs, especially when larger number of eggs were produced. Pupae are hardly missed, compared to eggs which tend to cluster together. Eggs and worker pupae might have been counted more than once, as development lasts a median of 8 and 18 days for eggs and worker pupae, respectively. Finally, three colonies (10 worker treatment) were not considered in the analysis as they were accidentally killed.

To examine the worker aging patterns, 40 focal worker pupae were set up in individual colonies with 10 or 20 marked workers. These two treatments were selected, as no significant differences were observed between the 20 workers and 30 workers colonies in terms of queen productivity. Marking was done by clipping the tarsae of the middle right leg. The colony was set up with brood (5 larvae in the 10 workers colonies, and 10 in the 20 workers colonies), and two wingless queens to avoid a queenless period. The number of marked workers, queens and larvae was standardized weekly to the assigned treatment, and newly produced pupae produced were removed. Dead marked workers were replaced with fresh worker pupae and clipped one or two days after eclosion to avoid confusion with the focal worker.

### Offspring investment

Freshly eclosed adult workers were sampled monthly for head width measurements (from the 3rd to 6th month of the queen’s life, and up to five workers depending on availability). Workers were dried, pinned, and blindly measured using a Keyence Microscope 200X. A single worker was chosen randomly and measured 10 times to obtain a proxy for measurement error (Mean = 383.61 μm, standard deviation=5.05 μm).

### Statistical tests

To test for significant differences between treatments, we used generalized linear mixed effects models within the R package glmmTMB (R version 3.5.2, (Pinheiro et al. 2011)) and a negative binomial distribution for count data. If the count data and caste investment ratios were log transformed, a Gaussian family distribution was used. The dependent variable was analyzed as a function of the fixed effects: treatment (Number of workers as a factor), and random effects: stock nest and box of origin, box of set up, set up date. All models were also graphically checked for consistency and model diagnostics were performed using the DHARMA package (R version 0.3.3.0, (Hartig 2020)). To test for differences in head width size, we used the average of the head width measurements of the workers per time point (each month). Predictions of the data were visualized using the loess method with the geom_smooth function and default span (ggplot2 v.3.3.2). Data is available as external database S1, and the R-script is filed under database S2.

---

**Fig. S1.**
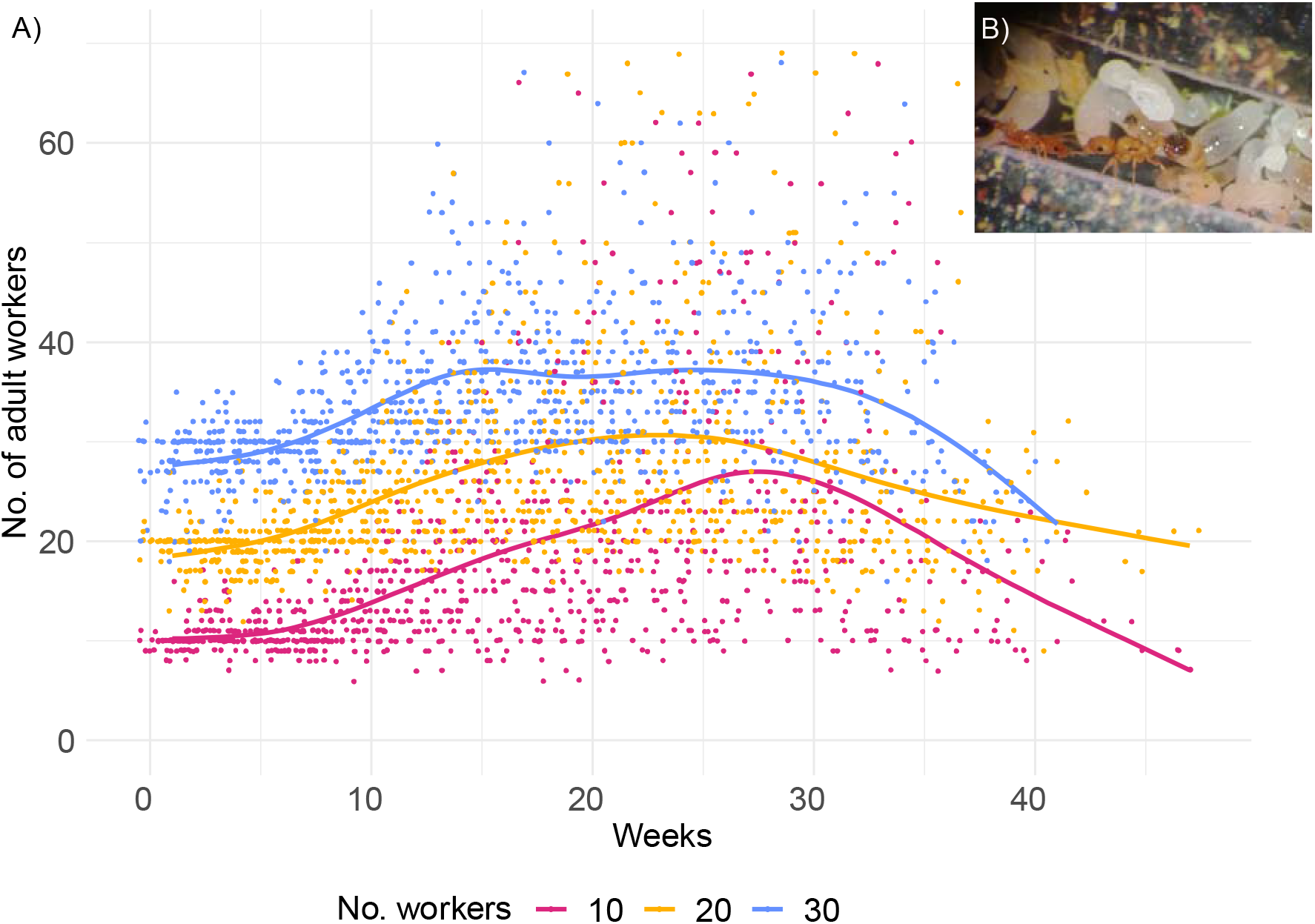
A) Adult ant workers present over the lifetime of the queens as a result of the colony size manipulation, B) *C. obscurior* ant queens, workers and colonies were monitored and standardized according to treatment (colony size) after the weekly count.

**Fig. S2.**
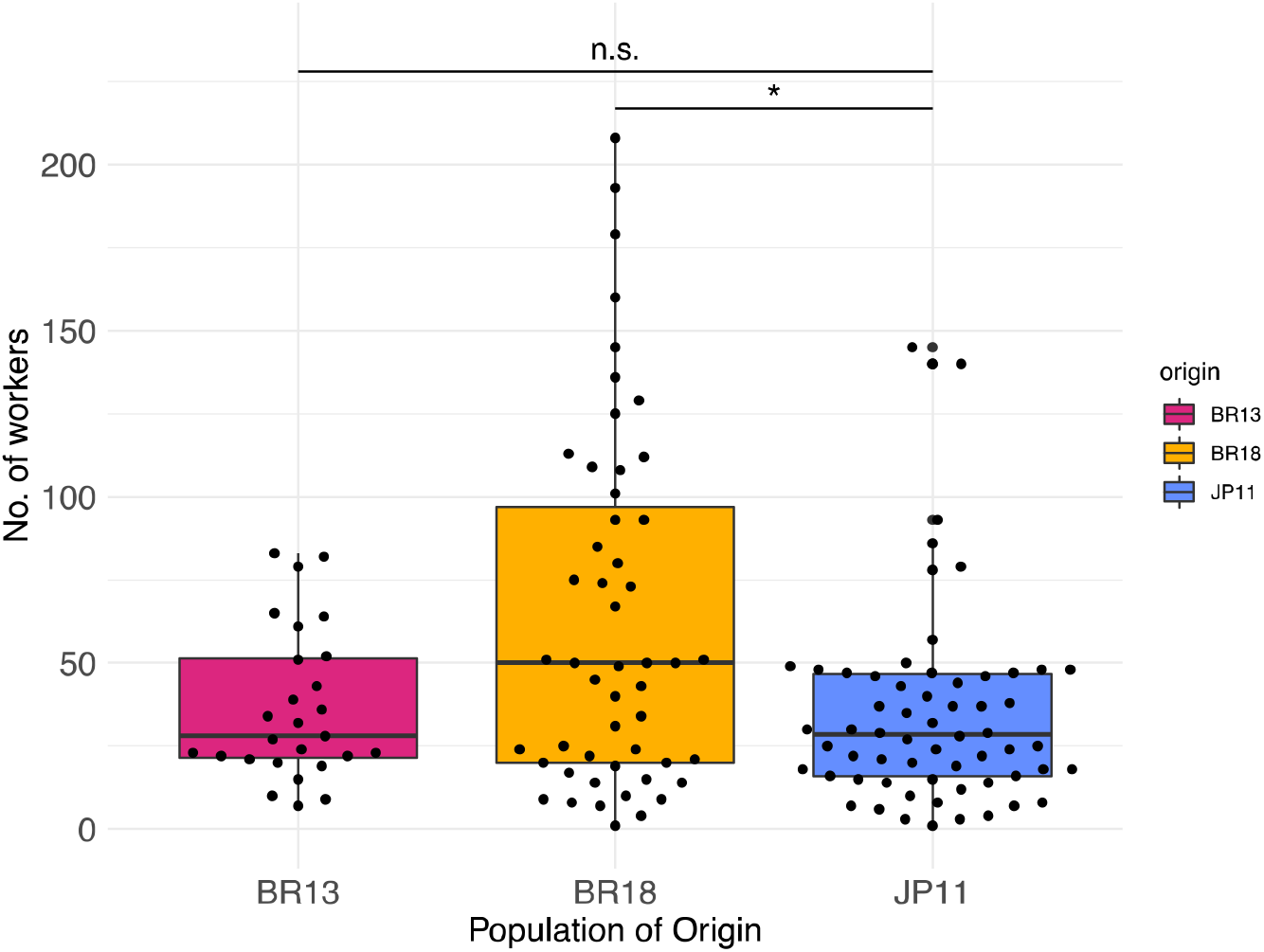
Worker numbers of colonies collected in 2013 (n=27) and 2018 (n=52) in Brazil, and in 2011 in Japan (n=62). One to 60 queens were found per colony. The Brazilian populations “BR13”, with a minimum = 7, lower-hinge = 21.5, median = 28, upper-hinge = 51.5, and maximum = 83, and “BR18”: minimum = 1, lower-hinge = 20, median = 20, upper-hinge = 104.5, and maximum = 244, and the Japanese population “JP11”: minimum = 1, lower-hinge = 16, median = 28.5, upper-hinge = 47, and maximum = 145. Colony size differed between populations (Kruskal-Wallis test, X^2^=8.34, p < 0.05), specifically among JP11 and BR18 (Post-hoc Kruskal-Nemenyi test, p < 0.05). Significant differences are given with * for p<0.05.

**Fig. S3.**
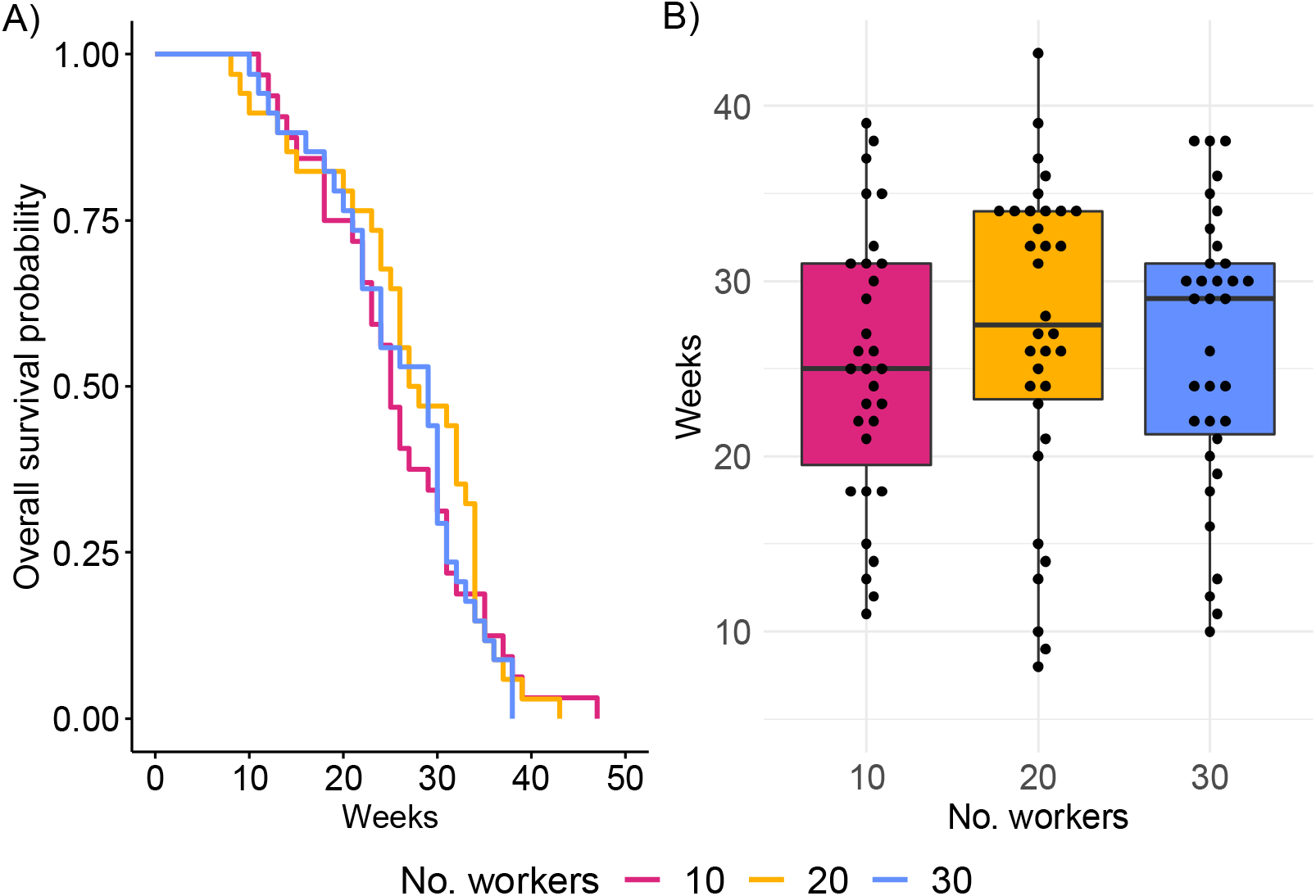
Queen longevity A) survival curve based on Cox proportional 335 hazard regression 336 model, and B) Queen lifespan in weeks.

**Fig. S4.**
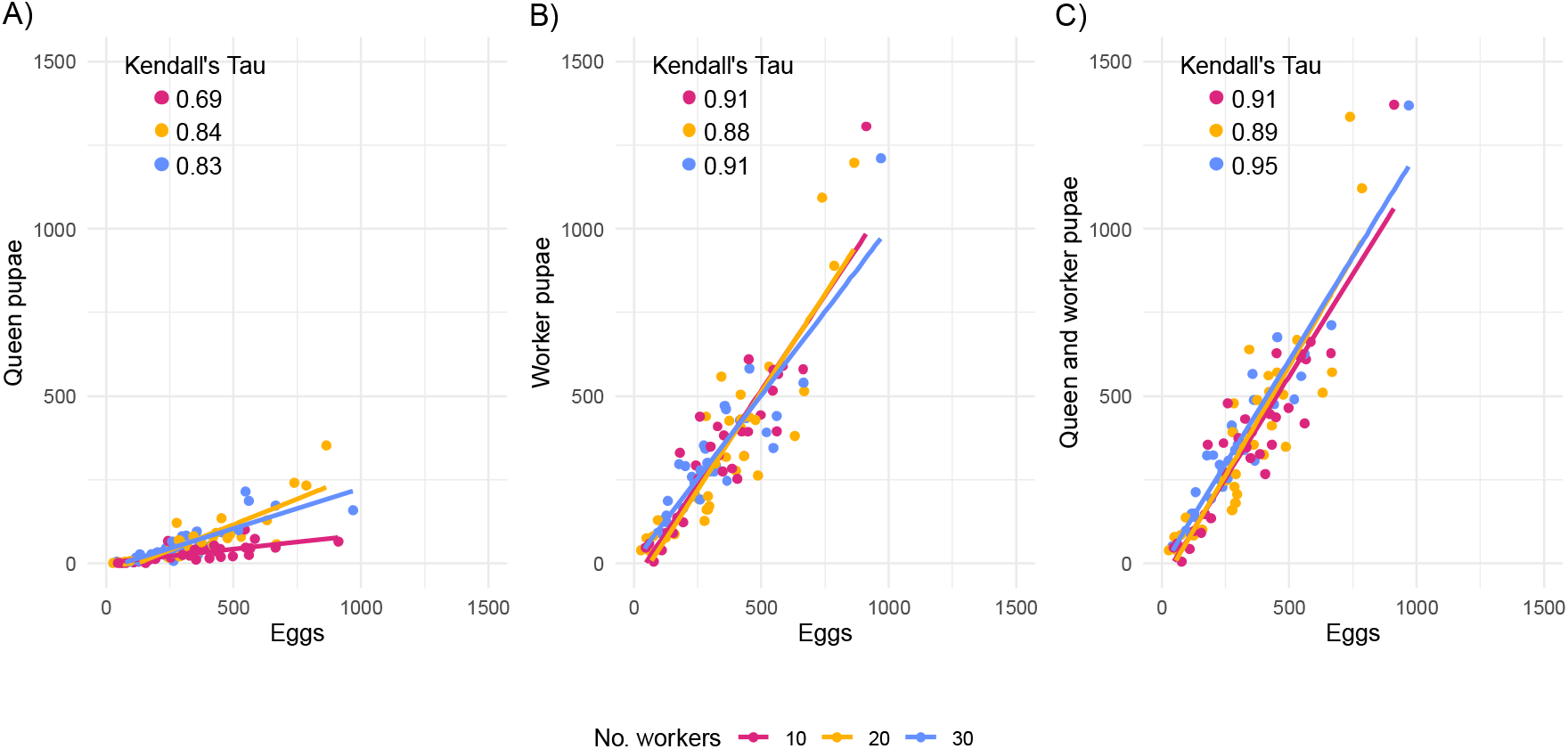
Correlation between counted eggs and A) Queen pupae, B) Worker pupae, and C) the sum of queen and worker pupae per queen. All correlations are statistically significant at p < 0.001 (Kendall’s rank correlation test).

## References

Beekman M, Lingeman R, Kleijne FM, Sabelis MW. 1998. Optimal timing of the production of sexuals in bumblebee colonies. Entomol. Exp. Appl. 88:147–154.

Boomsma JJ, Gawne R. 2018. Superorganismality and caste differentiation as points of no return: how the major evolutionary transitions were lost in translation. Biol. Rev. 93:28–54.

Bourke AFG. 2007. Kin selection and the evolutionary theory of aging. Annu. Rev. Ecol. Evol. Syst. 38:103–128.

Cohen AA. 2018. Aging across the tree of life: The importance of a comparative perspective for the use of animal models in aging. Biochim. Biophys. Acta - Mol. Basis Dis. 1864:2680–2689.

Corona M, Libbrecht R, Wheeler DE. 2016. Molecular mechanisms of phenotypic plasticity in social insects. Curr. Opin. Insect Sci. 13:55–60.

Flatt T, Partridge L. 2018. Horizons in the evolution of aging. BMC Biol. 16:1–13.

Giehr J, Heinze J, Schrempf A. 2017. Group demography affects ant colony performance and individual speed of queen and worker aging. BMC Evol. Biol. 17:1–9.

Haldane J. 1941. New Paths in Genetics. London◻: George Alien & Unwin, Ltd.

Hamilton WD. 1966. The moulding of senescence by natural selection. J. Theor. Biol. 12:12–45.

Harrison MC, Jaimes Nino LM, Rodrigues MA, Ryll J, Flatt T, Oettler J, Bornberg-bauer E. 2021. Gene co-expression network reveals highly conserved, well-regulated anti-ageing mechanisms in old ant queens. BioRxiv, https://doi.org/10.1101/2021.02.14.431190

Hartig F. 2020. DHARMa◻: Residual Diagnostics for Hierarchical (Multi-Level / Mixed) Regression Models. Available from: https://cran.r-project.org/package=DHARMa

Heinze J, Schrempf A. 2012. Terminal investment: Individual reproduction of ant queens increases with age. PLoS One 7:1–4.

Hughes KA, Reynolds RM. 2005. Evolutionary and mechanistic theories of aging. Annu. Rev. Entomol. 50:421–445.

Jones OR, Scheuerlein A, Salguero-Gómez R, Camarda CG, Schaible R, Casper BB, Dahlgren JP, Ehrlén J, García MB, Menges ES, et al. 2014. Diversity of ageing across the tree of life. Nature 505:169–173.

Keller L, Genoud M. 1997. Extraordinary lifespans in ants: a test of evolutionary theories of ageing. Nature 389:3–5.

Kirkwood TBL. 1977. Evolution of ageing. Nature 270:301–303.

Kirkwood TBL, Austad SN. 2000. Why do we age? Nature 408:233–238.

Kramer BH, Schaible R. 2013. Colony size explains the lifespan differences between queens and workers in eusocial Hymenoptera. Biol. J. Linn. Soc. 109:710–724.

Kramer BH, Schrempf A, Scheuerlein A, Heinze J. 2015. Ant colonies do not trade-off reproduction against maintenance. PLoS One 10:1–13.

López-Otín C, Blasco MA, Partridge L, Serrano M, Kroemer G. 2013. The hallmarks of aging. Cell 153:1194–1217.

Lucas E, Keller L. 2017. Explaining extraordinary lifespans◻: the proximate and ultimate causes of differential lifespan in social insects. In: Jones OR, Salguero-Gómez R, editors. The Evolution of Senescence in the tree of life. Shefferson. Cambridge: Cambridge University Press. p. 198–219.

Macevicz S, Oster G. 1976. Modeling social insect populations II: Optimal reproductive strategies in annual eusocial insect colonies. Behav. Ecol. Sociobiol. 1:265–282.

Maklakov AA, Chapman T. 2019. Evolution of ageing as a tangle of trade-offs: energy versus function. Proc. R. Soc. B Biol. Sci.:1–8.

Maklakov AA, Rowe L, Friberg U. 2015. Why organisms age: Evolution of senescence under positive pleiotropy? BioEssays 37:802–807.

Monroy Kuhn JM, Korb J. 2016. Editorial overview: Social insects: aging and the re-shaping of the fecundity/longevity trade-off with sociality. Curr. Opin. Insect Sci. 16:vii–x.

Moorad J, Promislow D, Silvertown J. 2020. Williams’ Intuition about Extrinsic Mortality Is Irrelevant. Trends Ecol. Evol. [Internet] 35:379. Available from: https://doi.org/10.1016/j.tree.2020.02.010

Oettler J, Schrempf A. 2016. Fitness and aging in *Cardiocondyla obscurior* ant queens. Curr. Opin. Insect Sci. 16:58–63.

Oster GF, Wilson EO. 1978. Caste and ecology in the social insects. Princeton, New Jersey, USA: Princeton University Press.

Pinheiro J, Bates D, Debroy S, Sarkar D. 2011. Linear and Nonlinear Mixed Effects Models.

Schrader L, Kim JW, Ence D, Zimin A, Klein A, Wyschetzki K, Weichselgartner T, Kemena C, Stökl J, Schultner E, et al. 2014. Transposable element islands facilitate adaptation to novel environments in an invasive species. Nat. Commun. 5:1–10.

Schrempf A, Cremer S, Heinze J. 2011. Social influence on age and reproduction: Reduced lifespan and fecundity in multi-queen ant colonies. J. Evol. Biol. 24:1455–1461.

Schrempf A, Giehr J, Röhrl R, Steigleder S, Heinze J. 2017. Royal Darwinian Demons: enforced changes in reproductive efforts do not affect the life expectancy of ant queens. Am. Nat. 189:436–442.

Schrempf A, Heinze J. 2006. Proximate mechanisms of male morph determination in the ant Cardiocondyla obscurior. Evol. Dev. 8:266–272.

Schrempf A, Heinze J, Cremer S. 2005. Sexual cooperation: mating increases longevity in ant queens. Curr. Biol. 15:267–270.

Toth AL, Sumner S, Jeanne RL. 2016. Patterns of longevity across a sociality gradient in vespid wasps. Curr. Opin. Insect Sci. 16:28–35.

